# Explainable machine learning reveals evolutionary signals in Influenza hemagglutinin

**DOI:** 10.1101/2025.09.21.677610

**Authors:** Austin G. Meyer

## Abstract

Identifying amino acid changes that lead to phenotypic change is a central problem that is critical to viral surveillance. Common metrics used to measure protein evolution like site-wise evolutionary rates and entropy are truly measures of variability rather than phenotypic importance. Here I show that supervised, explainable machine learning models provide a complementary approach that could serve to date and classify sequences, identify important mutations for host adaptation, and control directly for confounding covariates like sampling date and geography of origin. I curated 39,121 hemagglutinin (H3) protein sequences from GISAID with passage annotations and associated sample metadata to create models of sequence change. Gradient boosted decision trees were trained with encoded amino acids plus latitude, longitude, and date; SHAP values quantified site importance. The passage classifier achieved 81% overall accuracy (balanced accuracy = 0.77), distinguishing egg grown from unpassaged isolates with nearly 90% recall, and recovering known and novel adaptive substitutions. A separate regressor, trained solely on unpassaged sequences, predicted sample collection date with ***R***^***2***^ **= 0.98** and a mean absolute error of 74.5 days. Crucially, the sites identified as most important by the models showed a strong enrichment for experimentally validated antigenic sites, with the passage model ranking these functionally critical residues far more effectively than traditional evolutionary metrics. Across both tasks, correlations between SHAP values and standard evolutionary metrics were strong (**0.63 *≤ ρ ≤* 0.9**), indicating a strong connection between importance and variability depending on model specification. These results demonstrate that explainable machine learning can reveal important substitutions, deliver tree free molecular dating, and may transform passage metadata from a nuisance into an experimental probe.

## Introduction

Advances in sequencing have enabled real-time tracking of viral genetic change across scales, from global transmission to within-host evolution [1]. Many emerging pathogens, including pandemic influenza and HIV, originate from animal reservoirs and must rapidly adapt to new hosts [2]. Understanding which viral mutations confer fitness advantages, for example enabling immune escape or altering tropism, is critical for the development of new vaccines, vaccine strain updates, and outbreak control.

Classic studies of influenza A virus established that its evolution involves two different modes: the first involves gradual genetic drift while the second is characterized by punctuated antigenic “cluster” transitions. Early molecular analyses of influenza’s receptor binding and fusion protein, hemagglutinin (HA), identified specific codons under positive selection across influenza seasons [3], and sequence-based clustering showed H3N2 isolates grouping into discrete clades over time instead of forming a smooth continuum [4, 5]. Such epochal evolution may be driven primarily by immune selection [6, 7], and experimental studies have shown that even single mutations near the receptor-binding site were responsible for most H3N2 cluster transitions over the last several decades [8]. Thus, prospectively identifying such sites from sequences alone would significantly improve surveillance efforts. Identifying adaptive mutations from genetic data typically relies on site-wise metrics of variability and selection, such as Shannon entropy and the nonsynonymous/synonymous rate ratio (*dN/dS*) [9–12]. Traditionally, sites with *dN/dS >* 1 are interpreted as candidates for diversifying selection. However, these metrics are generally measures of relative variability, not necessarily of functional importance, and they face several challenges. First, *dN/dS* was derived for long-diverged lineages with fixed sequence differences and can behave unpredictably when applied to closely related sequences from an outbreak, such as during the 2009 H1N1 pandemic, the West African Ebola 2014 outbreak, or the early spread of SARS-CoV-2 [13–16]. Second, sites inferred to be under positive selection do not always correspond to known functional or antigenic sites [11]. Third, these metrics are easily confounded by technical artifacts and opposing evolutionary constraints. For example, a major source of spurious signals is *in vitro* sequence passaging for viral amplification, which introduces substrate-specific adaptations that can create false impressions of positive selection in nature [17].

To overcome these limitations, researchers have integrated additional context into evolutionary models. Protein structural features, such as solvent accessibility, weighted contact number, and proximity to functional sites (enzymatic sites or protein-protein interaction surfaces), can explain a fraction of evolutionary rate variation and improve inference in influenza and other viruses like HIV [11, 18–25]. However, these structural and rate-based metrics do not fully account for other biological factors that can strongly constrain amino acid substitution like glycosylation or epistasis that also modulate which residues evolve [26, 27]. To address these issues, there are many more advanced computational approaches that have been developed. Physics-based fitness models combining stability and activity have improved integration of multiple constraints [24]. Mechanistic fitness models have successfully predicted future dominant H3N2 clades [28]. Phylogeny-integrated frameworks have addressed the need for better strain forecasts [29], and deep mutational scanning experiments provide better understanding of the empirical fitness landscapes [30]. More recently, the flexibility of machine learning (ML) has been applied to predict upcoming influenza strains [31, 32] and to model protein evolution using large language models [33–38]. A key advantage of ML is the ability to integrate diverse data types—such as sequence, geography, and time—into a single framework. Unfortunately, the majority of these powerful ML models remain “black boxes”, and even when interpretation is possible, it is rarely implemented [32, 39].

However, in the context of public health, merely predicting an outcome is often insufficient in practice; understanding the underlying biological drivers is critical for building trust and making informed decisions. Explainable machine learning provides a path forward by allowing us to probe the logic of complex models [40, 41]. One of the most robust post-hoc explanation techniques is SHAP (SHapley Additive exPlanations), which assigns each feature a precise, theoretically grounded importance value for any given prediction [42]. By quantifying the marginal contribution of each amino acid, geographic coordinate, or time point, SHAP allows for a reliable assessment of what the model has learned.

This study tests the hypothesis that a feature’s (i.e., amino acid/site combination) predictive importance for a given phenotype can be a more direct and robust measure of its functional relevance than its site-wise variability. To this end, I present an explainable LightGBM framework that combines HA sequence data with geography, sampling date, and explicit passage-history labels. I demonstrate that this approach not only correctly classifies passage type and delivers accurate tree-free molecular dating, but also that its measure of feature importance aligns more closely with experimental data than do traditional metrics. Specifically, SHAP values rank experimentally validated antigenic sites far more effectively than do entropy, LEISR, or dN/dS. Furthermore, global SHAP rankings can identify relatively low-variability, phenotype-defining residues whose functional importance is confirmed by clear temporal frequency sweeps in the population data. By controlling for temporal, geographic, and laboratory dynamics within a single predictive architecture, this work converts passage metadata from a nuisance into an evolutionary probe and shows that supervised, interpretable machine learning can rival or surpass classical metrics for residue prioritization.

## 1 Results

### 1.1 Model performance on passage and dating tasks

I first trained two separate LightGBM models. One each on two prediction targets that together span the experimental and temporal co-variates of the dataset: (i) laboratory-passage history of each isolate and (ii) calendar date of collection for unpassaged sequences. After all quality filters (see Methods) the passage dataset comprised 39,121 H3 HA sequences annotated as Egg, MDCK, SIAT-MDCK, Monkey Kidney, or Unpassaged; the exceedingly rare Vero label (one sequence) was discarded. Splitting yielded 33,804 training and 8,452 heldout test sequences with nearly identical class proportions. Using the optimized hyperparameters recovered by Bayesian search, the classifier attained an overall test accuracy of 0.81 (balanced accuracy 0.77). The normalized confusion matrix (Fig. 1) shows that biologically distant classes are readily separated: egg-passaged isolates are predicted correctly in 90 % of cases and never mislabeled as Unpassaged, whereas unpassaged sequences achieve 94 % recall with negligible confusion for egg passaging. Errors concentrate among the three mammalian cell lines. In particular, it is difficult to distinguish Monkey and SIAT-MDCK passaging from unpassaged sequences. This is expected based on the biological similarity of the cell lines.

**Fig. 1:**
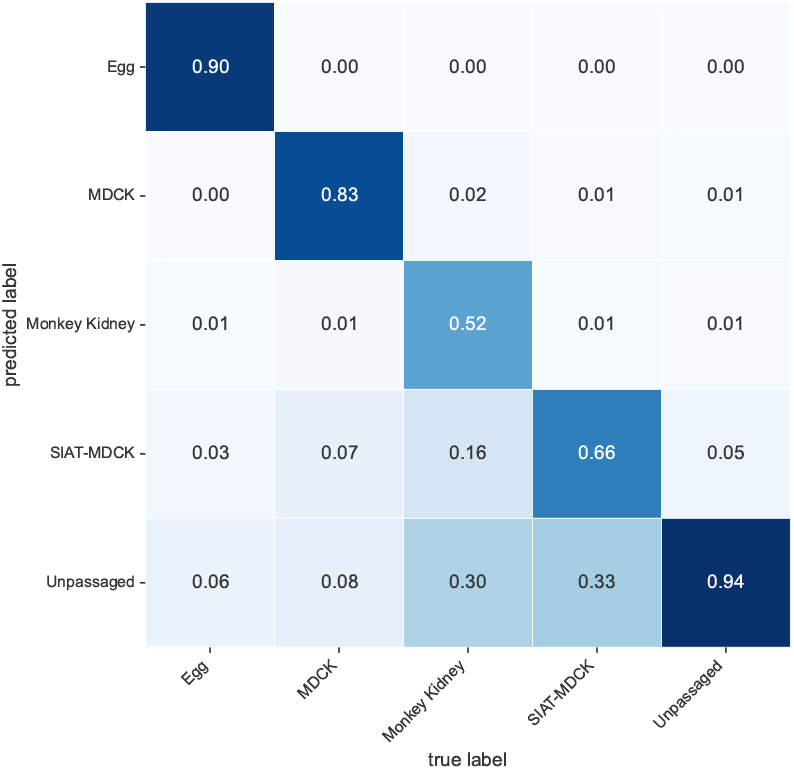
Classification performance. Normalized confusion matrix for six passage categories (test set); overall accuracy = 0.81, balanced accuracy = 0.77. Egg and unpassaged isolates are almost perfectly distinguished.

For molecular dating, I restricted the regression task to the 20,109 unpassaged sequences to avoid substrate-specific temporal signals. The data were split 80/20 (16,087 train, 4,022 test) by random sampling. Model inputs consisted solely of one-hot sequence features, latitude, and longitude; no explicit clock information or phylogenetic structure was provided. Predicted versus true collection dates exhibit a tight linear relation (Fig. 2), with *R*^2^ = 0.98 on the test set and a mean absolute error (MAE) of 74.5 days. For context, a baseline that uses only geographic covariates yields *R*^2^ = 0.13, whereas a “sequence-only” model already reaches *R*^2^ = 0.98– indicating that temporal signal is encoded predominantly in the amino-acid pattern. Importantly, the LightGBM predictions on the test set also correlate strongly with classical root-to-tip genetic distances (*r* = 0.94), yet are obtained without an explicit tree or molecular-clock fit.

**Fig. 2:**
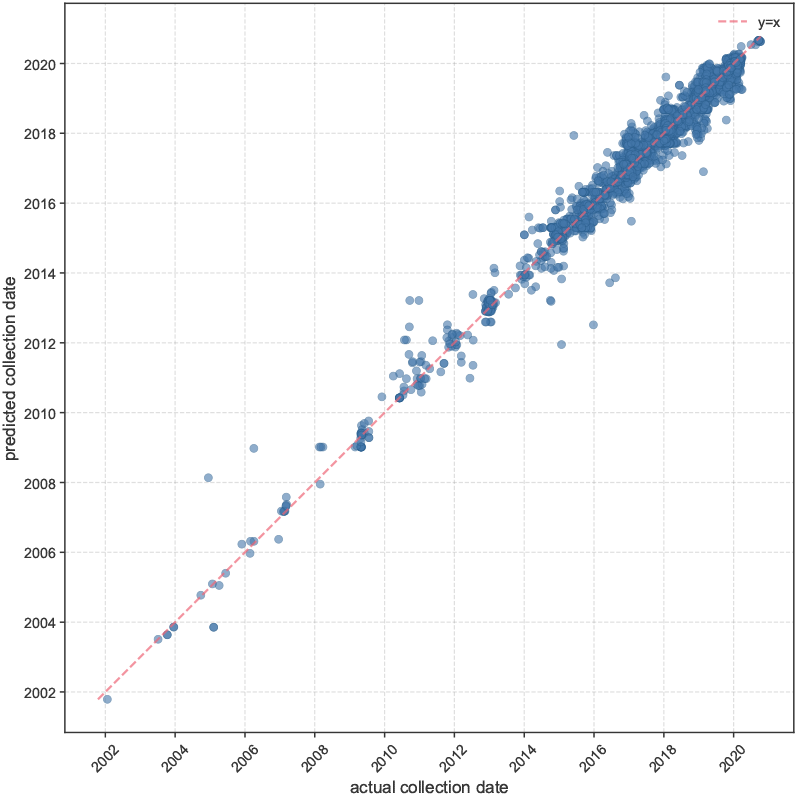
Tip-date regression. LightGBM predictions versus actual collection dates for 4022 unpassaged test sequences (*R*^2^ = 0.98, MAE = 74.5 d). Dashed line indicates *y* = *x*.

Together these results demonstrate that a single gradient-boosting architecture can (i) learn the discrete, non-phylogenetic lab-passage labels with good accuracy and (ii) solve the continuous tip-dating problem with near-molecular-clock precision. The strong performance of the models across tasks justifies interrogating the model to understand which specific residues drive passage differentiation and which encode temporal signal.

### 1.2 Global SHAP rankings identify key residues

To understand which features were most influential in the models, I employed SHAP values to quantify the contribution of each input feature (individual amino acid states at each position, latitude, longitude, and time for the passage model) to the predictions. For the passage classification model, the global SHAP analysis aggregates the importance of features across all predictions, revealing which amino acid substitutions, or geographical features, most strongly differentiate passage histories. Being continuous variables, geographical features (latitude and longitude) and collection time contribute strongly to passage classification as three of the four most important individual features in the model (Fig. 3). However, I found that after controlling for location and time, mean absolute SHAP values highlight specific positional amino acids that are strongly predictive for certain passage types (Fig. 3). For instance, the presence of threonine at protein site 160 (160T, corresponding to gene site 176) is a dominant feature for identifying egg-passaged sequences. Similarly, asparagine at protein site 144 (144N, gene site 160) emerges as a key predictor for monkey kidney-passaged isolates, and leucine at protein site 3 (3L, gene site 19) is a strong predictor of MDCK passage. Additionally, several amino acid-sites among the top 15 in Fig. 3 are known passage adaptation markers [17] based on analysis of sequence variability (to be discussed in section 1.5). However, many others among the top 15 are physically near to, but not specifically identified as, important sites in prior analysis of sequence variation, suggesting variation alone is insufficient to identify passage adaptation.

**Fig. 3:**
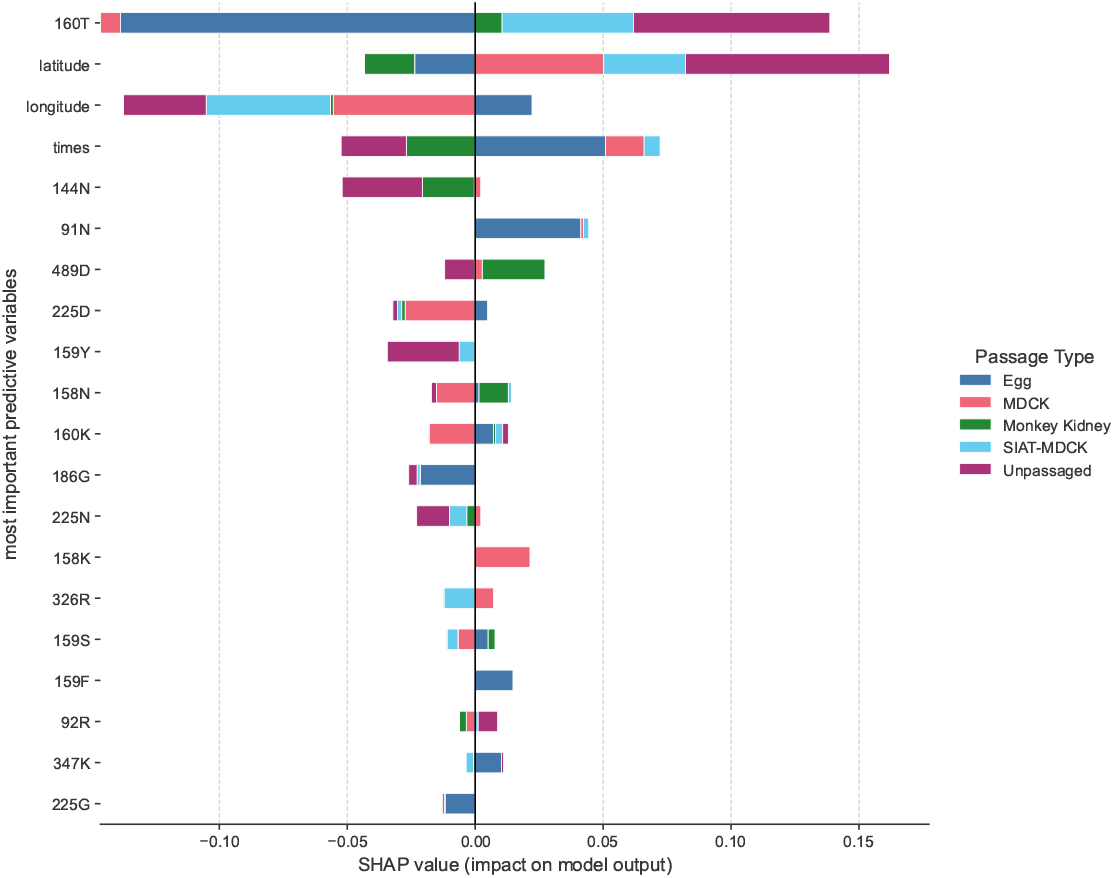
Passage model feature importance. Normalized SHAP values summed by residue and coloured by passage class. Features are ranked by sum for each site. Thus, the wider a bar is the more important it is for passage prediction. Negative values mean the feature was negatively associated with that passage type and positive values mean the feature was positively associated with that passage type. For example, the most important feature 160T is negatively associated with egg passaging and positively associated with unpassaging; thus, it is generally a wild type sequence. By contrast, 159F is positively associated with egg passaging implying that it is primarily an egg adaptive mutation.

In the date regression model, which was trained exclusively on unpassaged sequences and used only sequence and geographical data (latitude and longitude) as input, site-level SHAP values identify the amino acid positions most critical for predicting the sampling date. Aggregating SHAP values for all amino acids observed at each site reveals the overall importance of each position in the HA protein for temporal prediction. The analysis shows that protein sites 142, 3, and 144 (corresponding to gene sites 158, 19, and 160 respectively) are among the most influential predictors of sequence collection date (Fig. 4). The high importance of these sites suggests they have undergone consistent temporal changes that the model leverages for dating. For example, as will be shown in section 1.4, protein site 142 exhibits a clear R→ G→ R→ K→ G substitution pattern over time, and site 3 shows an L→I shift, both of which are captured by their high SHAP values. This demonstrates the model’s capacity to identify residues undergoing evolutionary changes that correlate strongly with time, without any explicit phylogenetic input.

**Fig. 4:**
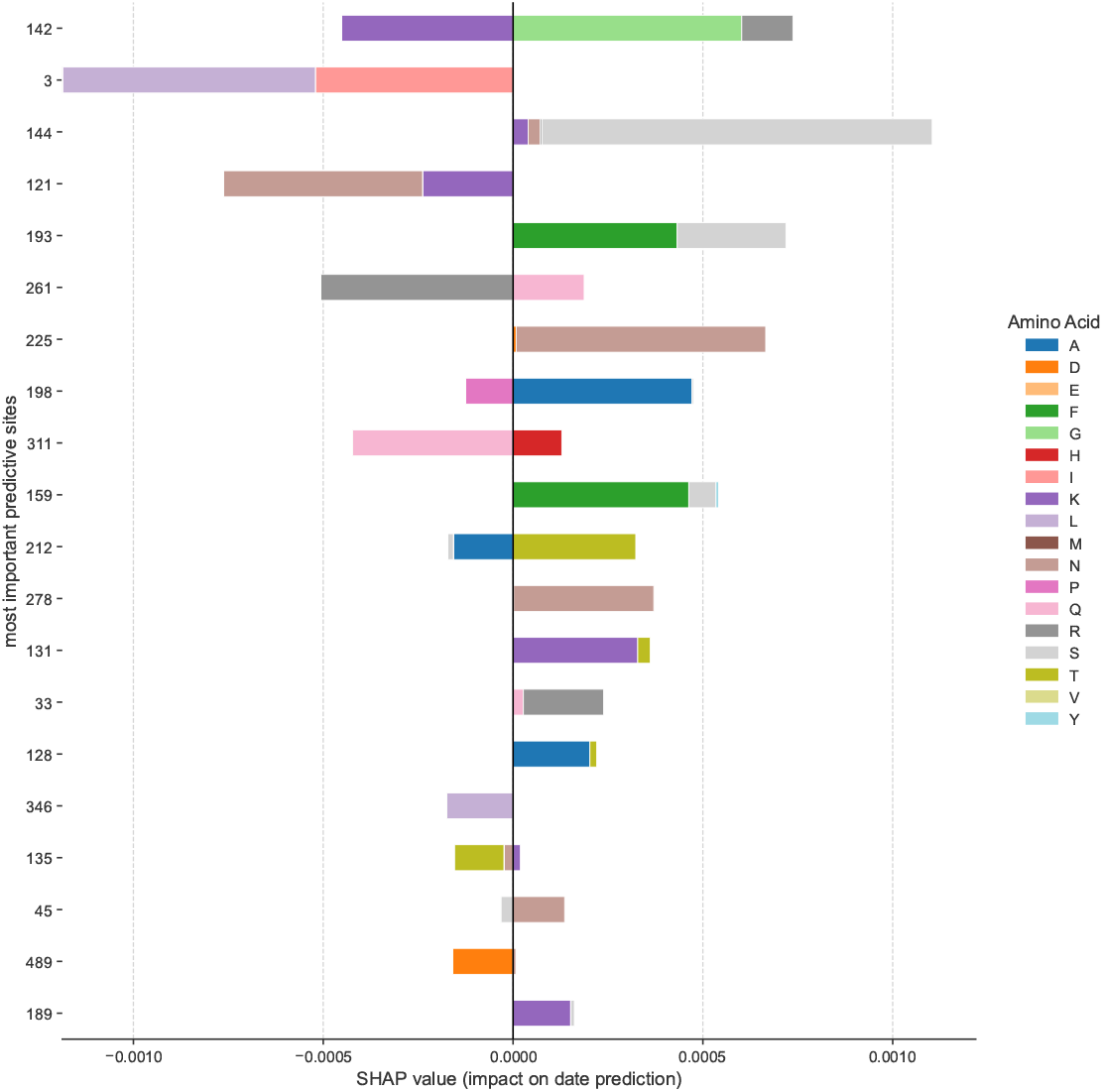
Site-level importance in the tip-date model. Bars show summed SHAP contribution per HA position. Features are ranked by sum for each site. Thus, the wider a bar is the more important it is for date prediction. Sites 142, 3, and 144 emerge as dominant predictors of sampling time, motivating the frequency-trajectory plots in Fig. 9.

### 1.3 SHAP probes site importance separate from variability metrics

While global SHAP rankings identify residues critical for model predictions (Figs. 3 and 4), it is important to understand how this measure of site specific importance relates to traditional metrics of sequence variability and evolutionary pressure. Classical approaches such as Shannon entropy, relative evolutionary rate (LEISR), and site wise nonsynonymous/synonymous substitution rate ratios (*dN/dS*) primarily measure sequence diversity or specific modes of selection [11, 14]. However, these methods may overlook sites that are functionally important but relatively conserved.

To benchmark SHAP against these traditional metrics, I assessed the Spearman correlations among them (Fig. 5). For the passage model, SHAP values correlated strongly with Shannon entropy (*ρ* = 0.90; Fig. 6), LEISR (*ρ* = 0.83; Fig. 7), and dN/dS (*ρ* = 0.77; Fig. 8). The correlations were consistently positive but less pronounced for the tip date regression model. For this model, SHAP values correlated with entropy at *ρ* = 0.76 (Fig. 6), with LEISR at *ρ* = 0.70 (Fig. 7), and with dN/dS at *ρ* = 0.63 (Fig. 8). These results show that while predictive importance is related to sequence variability, the strength of this relationship depends on the biological question being asked. The moderate correlations, particularly for the date model, indicate that SHAP captures a dimension of functional importance not fully described by variability metrics alone.

**Fig. 5:**
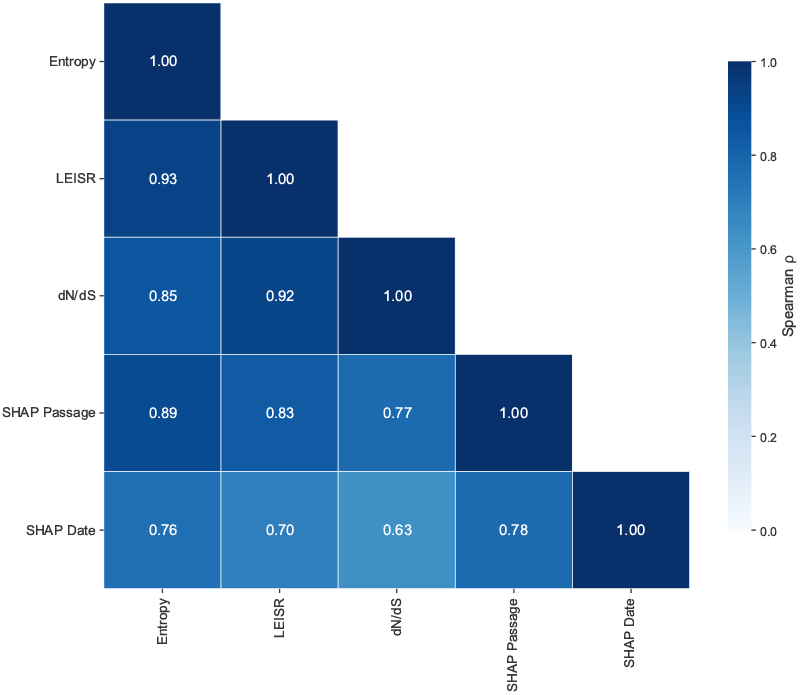
Spearman correlation heatmap. The strength of the Spearman correlation between the metrics studied in this paper. There is a moderate-to-strong correlation among all of the metrics depending on the model structure of the LightGBM model.

**Fig. 6:**
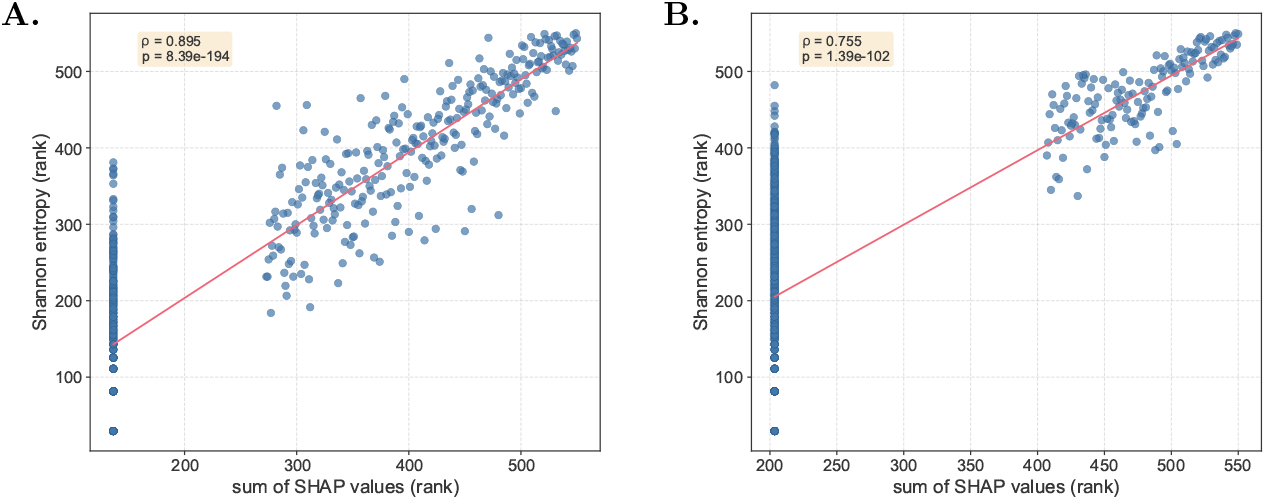
SHAP versus sequence entropy. Correlation between summed SHAP values and Shannon entropy for **A**. passage classification (*ρ* = 0.90) and **B**. tip-date regression (*ρ* = 0.76). In general, there is a moderate-to-strong correlation between entropy and SHAP. However, there are many sites with relatively high entropy that offer no value in predicting the evolution of the protein.

**Fig. 7:**
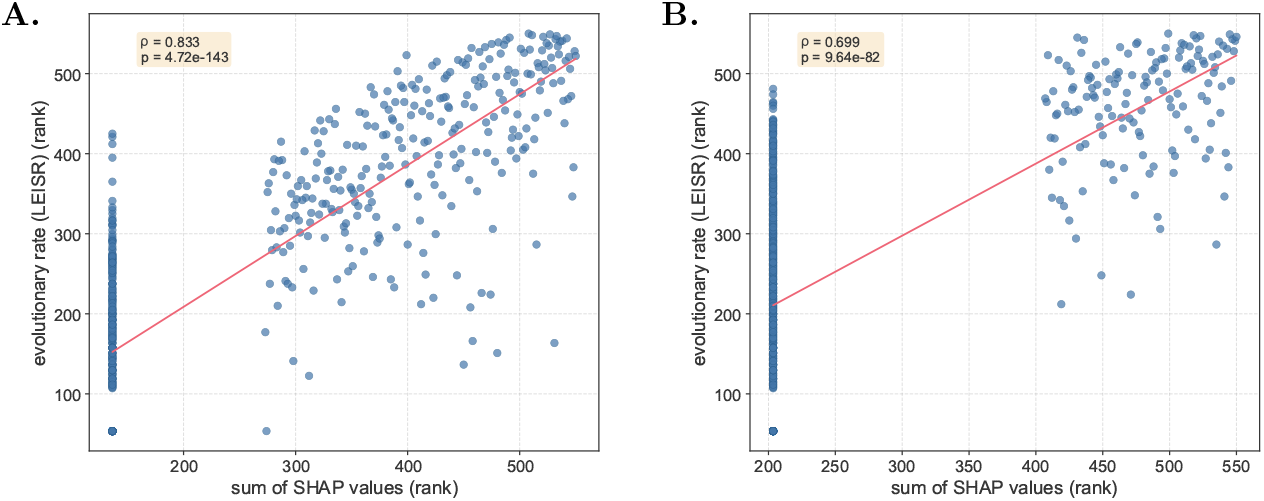
SHAP versus relative evolution rate. **A**. Passage classifier (*ρ* = 0.83). **B**. Tip-date regressor (*ρ* = 0.69). As with entropy, there is a moderate-to-strong correlation between relatively evolutionary rate and SHAP. Again though, there are many sites with high LEISR that are of no value in predicting the evolution of the protein.

**Fig. 8:**
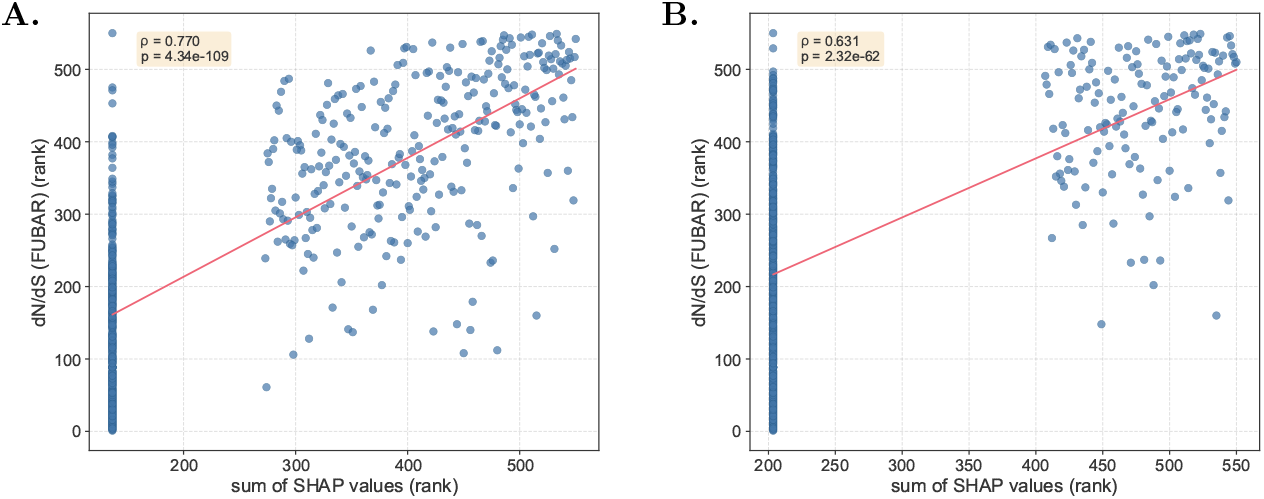
SHAP versus dN/ds. **A**. Passage classifier (*ρ* = 0.77). **B**. Tip-date regressor (*ρ* = 0.63). As with entropy and LEISR, there is a moderate-to-strong correlation between relatively evolutionary rate and SHAP. Yet, we find a significant number of high dN/dS sites that offer no value in predicting the evolution of the protein.

### 1.4 Temporal sweeps confirm functional importance of top sites

To further validate the biological relevance of sites identified as important by SHAP, particularly in the context of temporal prediction, I examined the amino acid frequency trajectories of top-ranking amino acid sites over the sampling period. The capacity of the tip-date regression model to accurately predict collection dates (Fig. 2) relies on its ability to detect consistent evolutionary changes. A temporal sweep, where one amino acid is replaced by another across the population over time, represent a strong evolutionary signal. Note, these sweeps are examined solely in the unpassaged sequence set.

I focused on two illustrative examples, protein site 142 and site 3, which were among the most influential predictors of sampling date according to their aggregated SHAP values (Fig. 4). At site 142 in terms of primary amino acid at the site, an arginine (R) changed to glycine (G) then briefly back to arginine (R) before moving to lysine (K) and back to glycine (G) before fixing (R→ G→ R→ K→ G); this swept through the H3N2 population between approximately 2012 and 2020 (Fig. 9, left panel). Similarly, site 3 experienced a clear leucine (L) to isoleucine (I) transition (L→ I) that started in 2012 and fixed around 2018 (Fig. 9, right panel).

**Fig. 9:**
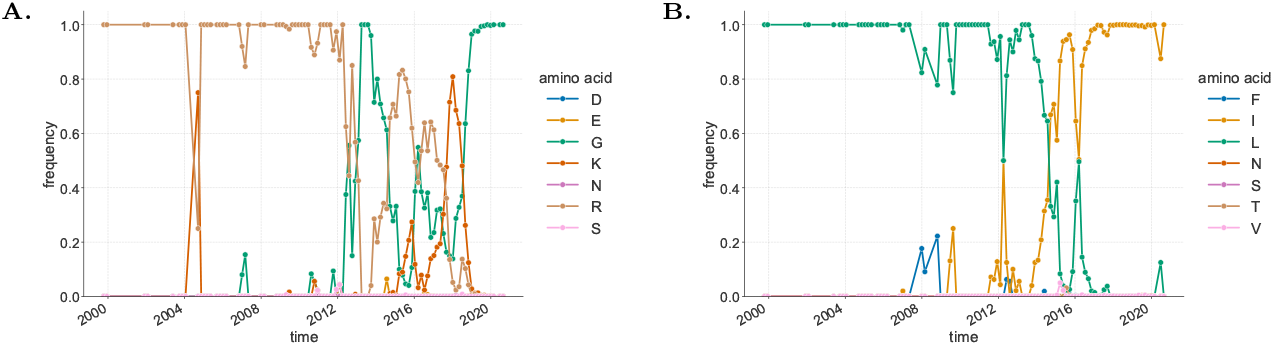
Sweeps of SHAP-ranked residues. Amino-acid frequency trajectories for **A**. protein site 142 (R→ G→ R→ K→ G→ sweep, 2012–2020) and **B**. protein site 3 (L→I sweep, 2012–2018). Both residues rank among the most influential for sampling-date prediction yet have modest entropy, validating their biological relevance.

These observed frequency changes provide evidence for the importance of these SHAP-ranked residues. For reference, protein site 142 has a relative evolutionary rate greater than 6, which is among the highest in the protein. However, among the other top 10 sites according to summed SHAP value, site 311 (the 9th most important site for date regression) has a LEISR value of only 1.3. Of the remaining top 10, protein site 3 has a *LEISR* = 4.9, protein site 144 has a *LEISR* = 4.2, protein site 121 has a *LEISR* = 5.2, protein site 193 has a *LEISR* = 2.7, protein site 261 has a *LEISR* = 7.4, protein site 241 has a *LEISR* = 1.1, protein site 198 has a *LEISR* = 4.6, and protein site 159 has a *LEISR* = 0.8. Thus, despite these sites exhibiting significantly different LEISR values (as discussed in Section 1.3), they are strong predictors of sampling time. The LightGBM model, as interpreted by SHAP, successfully identifies these residues as critical for dating sequences. This demonstrates that the explainable machine learning framework can pinpoint residues undergoing significant evolutionary transitions that are directly relevant to the temporal progression of the virus, and therefore confirm the model’s ability to capture biologically meaningful evolutionary signals without explicit phylogenetic information.

### 1.5 Comparison to experimentally determined antigenic sites

To benchmark the predictive importance derived from SHAP against classical evolutionary metrics, I assessed the rankings of seven experimentally validated antigenic sites in H3 hemagglutinin previously identified by Koel et al. [8]. These sites (145, 155, 156, 158, 159, 189, and 193) are known to be critical for the antigenic evolution of the virus. We ranked all sites in the HA protein according to each metric—SHAP (passage and date models), dN/dS, LEISR, and entropy—and then examined the specific ranks of these seven key sites (Table 1).

**Table 1:** Antigenic Site Rankings. Importance ranks for the seven key antigenic sites identified by Koel et al. [8]. The table shows each site’s rank out of all protein sites for five different metrics, where a lower number indicates higher importance. Metrics include SHAP values from the passage (*SHAP*_*passage*_) and date (*SHAP*_*date*_) models, dN/dS, LEISR, and Shannon entropy.

| Protein Site | Gene Site | $SHAP_{passage}$ | $SHAP_{date}$ | dN/dS | $LEISR$ | Entropy |
| --- | --- | --- | --- | --- | --- | --- |
| 145 | 161 | 28 | 66 | 95 | 44 | 30 |
| 155 | 171 | 82 | 246 | 281 | 325 | 69 |
| 156 | 172 | 102 | 122 | 266 | 198 | 60 |
| 158 | 174 | 4 | 28 | 66 | 79 | 44 |
| 159 | 175 | 3 | 8 | 117 | 204 | 3 |
| 189 | 205 | 11 | 19 | 120 | 113 | 42 |
| 193 | 209 | 16 | 9 | 18 | 60 | 15 |

The analysis revealed that SHAP-based importance, particularly from the passage model, prioritized the known antigenic sites more effectively than did the other metrics. The passage model’s SHAP values placed four of the seven antigenic sites within the top 20 ranks, including two in the top 10 (sites 159 and 158). In contrast, the classical metrics assigned substantially lower importance to these sites. Neither dN/dS nor LEISR ranked any of the seven antigenic sites in the top 10. Furthermore, among the antigenic sites, dN/dS indicated that only protein site 193 (gene site 209) was under positive selection with dN/dS = 1.4; no other sites are undergoing positive selection. Entropy performed better than the rate-based metrics but still only identified one site in the top 10 (site 159).

A summary of these findings highlights the difference in performance across the metrics (Table 2). The median rank for the seven antigenic sites was lowest for SHAP from the passage model (16), followed by SHAP from the date model (28) and Shannon entropy (42). The evolutionary rate metrics performed most poorly, yielding median ranks of 113 for LEISR and 117 for dN/dS. These results show that the supervised, phenotype-aware importance assigned by the SHAP framework aligns more closely with experimentally determined antigenic sites than do unsupervised measures of variability or evolutionary rate.

**Table 2:** Summary of Antigenic Site Rankings. Median and mean ranks for the seven antigenic sites listed in Table 1, summarized across the five metrics, and ordered by median rank. Lower values indicate that a metric assigns a higher average importance to the set of known antigenic sites.

| Metric | Median Rank | Mean Rank |
| --- | --- | --- |
| SHAP<sub>passage</sub> | 16 | 35.1 |
| SHAP<sub>date</sub> | 28 | 71.1 |
| Entropy | 42 | 37.6 |
| LEISR | 113 | 146.1 |
| dN/dS | 117 | 137.6 |

### 1.6 Structural location of SHAP-prioritised and antigenic sites

To understand the biophysical relevance of the sites identified as important by the passage model, I mapped their locations onto the crystal structure of H3 hemagglutinin in complex with its human receptor, sialic acid (Figure 10). The ten highest-ranking sites from the passage model’s SHAP analysis were visualised alongside the seven experimentally validated antigenic sites from Koel et al. [8].

**Fig. 10:**
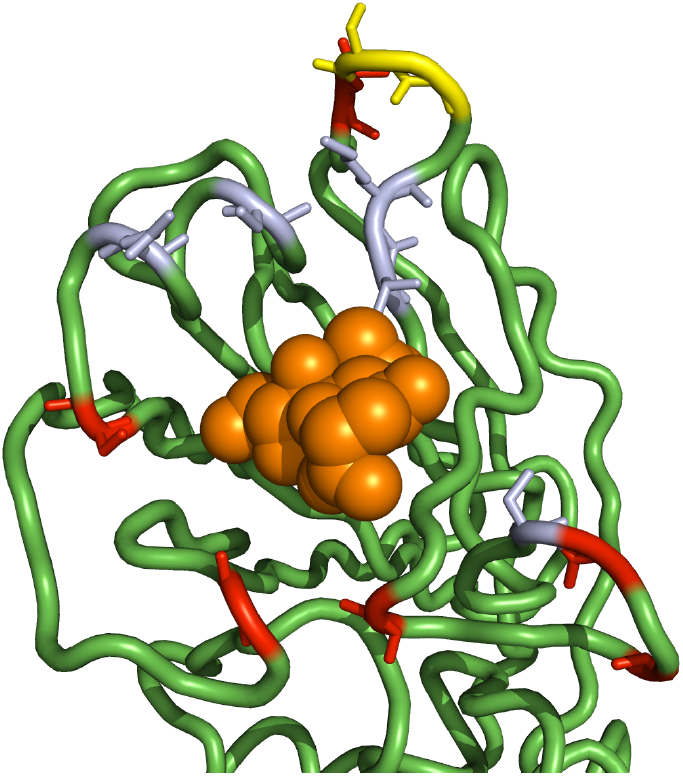
Structural location of important sites. The structure shows the cartoon representation of the H3 protein’s sialic acid binding site. Sialic acid is shown in orange. Green is the *α*-carbon backbone. Sites colored lavender are the antigenic sites identified by Koel et al. Sites colored red are among the top 10 most important sites according to SHAP for the passage model. Sites colored yellow are the two overlap sites between the antigenic sites and the top 10 shap sites.

The structural mapping reveals that the sites most predictive of passage history form a distinct cluster immediately adjacent to the sialic acid binding pocket. This proximity suggests their functional importance is directly related to receptor interaction, a key aspect of host adaptation. Notably, two of the top four SHAP-ranked sites, protein sites 158 and 159, are also members of the Koel et al. antigenic set, highlighting a clear intersection between predictive importance and known antigenicity. The remaining eight top-ten SHAP sites, while not in the Koel et al. set, are also located in the immediate vicinity of the receptor. This tight spatial clustering provides a strong biophysical rationale for their high predictive power in the passage model, as substitutions in this critical region are likely to alter binding characteristics.

## 2 Discussion

In this study, I have demonstrated that an explainable machine learning framework can effectively identify functionally important residues in influenza HA, offering a powerful complement to traditional evolutionary metrics. By training gradient-boosting models on a large dataset of H3N2 sequences with associated metadata, I have shown that predictive importance, as quantified by SHAP values, reveals biological signals that are often decoupled from measures of site-wise variability like Shannon entropy, LEISR, and dN/dS. The models not only performed with high accuracy on classification and regression tasks but, more importantly, provided a new lens through which to interpret the evolutionary pressures shaping the virus.

The passage classification model successfully learned the molecular signatures of laboratory adaptation. It distinguished egg-passaged isolates from unpassaged sequences with high recall (Figure 1), and the SHAP analysis identified key residues, such as 160T, that are strongly predictive of passage history (Figure 3) and were recently recapitulated by very detailed mechanistic studies [27]. The biophysical relevance of these findings is underscored by the structural location of the top-ranked sites. As shown in Figure 10, the most important residues for passage classification form a distinct cluster around the sialic acid receptor binding site, directly implicating this region in the adaptive changes that occur during laboratory processing. This result serves two purposes: first, it validates that the model is learning biologically meaningful features that align with known passage adaptations [17, 26, 27]; second, it demonstrates how passage metadata, often treated as a confounding variable to be filtered out, can be leveraged as an experimental probe. The model’s difficulty in distinguishing between certain mammalian cell lines (e.g., MDCK and SIAT-MDCK) is itself an informative finding, suggesting these environments exert similar, human-like selective pressures. In addition, this framework could be adapted to identify signatures of host-switching events, a critical task for pandemic preparedness [2].

Perhaps the most striking result is the performance of the tree-free tip-dating model. By using only sequence and geographic data, the LightGBM regressor predicted sampling dates with remarkable accuracy on a validation set (*R*^2^ = 0.98, MAE = 74.5 days), correlating strongly with traditional phylogenetic root-to-tip distance methods without the computational overhead of tree construction and molecular clock model fitting. This finding is particularly relevant given the known time-dependency of molecular rate estimates, which can complicate phylogenetic dating over short timescales [13, 14]. The SHAP analysis pinpointed sites 142, 3, and 144 as the most informative for this task (Figure 4). The validation of these sites through temporal frequency analysis, which revealed clear adaptive sweeps at sites 142 (R→ G→ R→ K→ G) and 3 (L→ I) (Figure 9), confirms that the model identified genuine, chronologically structured evolution. This provides a rapid, scalable alternative for dating large numbers of sequences in real-time genomic surveillance.

A central thesis of this work is that predictive importance is not synonymous with variability. This distinction is powerfully illustrated by comparing each metric’s ability to identify experimentally validated antigenic sites. While the correlations between site-wise SHAP values and metrics like entropy and LEISR were moderate to strong, SHAP proved far more effective at prioritizing functionally critical residues. The passage model’s SHAP values ranked the seven key antigenic sites from Koel et al. [8] with a median rank of 16, substantially outperforming entropy (median rank 42), LEISR (113), and dN/dS (117) (Table 2). Classical methods favor sites with high diversity or those under strong positive selection [7, 12]. However, a residue that is critical for a major antigenic shift or functional change may be under strong purifying selection before and after the event, exhibiting low overall variability [11]. This framework captures this by identifying sites whose state is highly predictive of a phenotype (e.g., passage type or date), regardless of their overall conservation profile. This moves beyond simply asking “which sites are variable?” to “which sites are informative for a specific biological outcome?”, a subtle but important distinction for understanding viral adaptation [8, 28].

This study has limitations. The analysis is confined to H3N2 HA, and the generalizability of these specific findings to other influenza subtypes or other viruses remains to be tested. The performance of any machine learning model is contingent on the quality and quantity of the training data, and is susceptible to sampling biases inherent in global surveillance data. While LightGBM is a powerful algorithm, other architectures, including deep learning models like transformers that have shown promise in protein modeling [33, 35], may capture different and important aspects of sequence evolution. Future work should focus on extending this explainable framework to other pathogens, integrating diverse data types such as serological or structural data, and comparing the feature attributions from different model architectures to build a more comprehensive picture of viral fitness landscapes.

In conclusion, this work establishes explainable machine learning with gradient-boosted decision trees as a valuable and practical tool for evolutionary virology. It provides a robust method for identifying phenotype-defining mutations that may be missed by classical approaches, offers a computationally efficient method for molecular dating, and provides a framework for converting experimental metadata into biological insight. By bridging the gap between predictive power and biological interpretability, this approach can enhance my ability to monitor viral evolution and inform public health strategies in an increasingly data-rich world.

## 3 Methods

### 3.1 Data Acquisition and Preprocessing

#### 3.1.1 Sequence Retrieval and Initial Filtering

Hemagglutinin protein sequences of influenza A(H3N2) virus, along with their corresponding metadata, were downloaded from the GISAID EpiFlu database (gisaid.org) [43]. The dataset spanned human isolates collected globally between 1993 and 2021 (though there were very few sequences available after the start of the COVID-19 pandemic in March of 2020). Initially, 72,690 sequences were retrieved. Metadata included isolate ID, collection date, location (country, region, city), and passage history among other things. Corresponding nucleotide sequences were also prepared for phylogenetic and evolutionary rate analyses.

Sequences were subjected to a series of filtering steps to ensure data quality for downstream analyses. I retained only sequences with complete passage history information, where passage details were not marked as ‘UnknownCell’. Isolates without a valid collection date or country information were excluded. Furthermore, only sequences possessing a complete HA protein sequence of 566 amino acids without any internal gaps or ambiguous characters (e.g., ‘X’, ‘-’) were kept. This comprehensive filtering resulted in a final dataset of 39,121 sequences. For site numbering, I distinguish between “gene site” (or “FASTA site”), referring to the 1-indexed position in the full gene present in the FASTA file, and “protein site”, which refers to the 1-indexed position in the 550 amino acid mature HA protein after cleavage of the N-terminal 16-amino-acid signal peptide. Thus, protein site *n* corresponds to gene site *n* + 16. All site numbers reported in figures and discussions of SHAP values refer to protein sites unless otherwise specified.

#### 3.1.2 Passage History Parsing

Passage history strings from the metadata were systematically parsed using the flupan Python library [17]. This process extracted up to four distinct passage events for each isolate. The parsed passage information was cached to avoid redundant processing in subsequent analyses. Based on the primary passage event (pass1), sequences were categorized. If a second passage event (pass2) was recorded and was not ‘UNPASSAGED’, the primary passage label was consolidated to ‘MULTI’ and all ‘MULTI’ sequences were removed from the analysis. Any sequence explicitly labeled ‘UNPASSAGED’ in any passage slot was assigned ‘UNPASSAGED’ as its final primary passage label. All passage labels were converted to uppercase for consistency (e.g., ‘EGG’, ‘MDCK’, ‘SIAT’, ‘MONKEYKIDNEY’, ‘UNPASSAGED’).

#### 3.1.3 Geographic Data Acquisition and Processing

The country of origin for each isolate was used to obtain geographical coordinates (latitude and longitude). Unique country names were geocoded using the Google Geocoding API via the geopy Python library. To minimize API calls and processing time, results were cached locally. For isolates where country-level geocoding failed or was ambiguous, latitude and longitude were treated as missing values. During feature preparation for machine learning models, missing latitude and longitude values were imputed using the median of the available coordinates in the training set.

#### 3.1.4 Sequence Encoding and Feature Preparation

For input into the machine learning models, HA protein sequences were converted into numerical features. Each of the 566 amino acid sites in the protein were one-hot encoded for each distinct amino acid appearing at each site in the alignment. This process creates a binary vector for each site/amino acid combination, where each position in the vector corresponds to a possible amino acid, and a ‘1’ indicates the presence of that amino acid at that site, with ‘0’s elsewhere. For example, if there are five different amino acids that appear at site 1 in the protein, the first amino acid site in the protein would take the first five positions in the one-hot vector. If site 2 in the protein had three different amino acids in the alignment, site 2 would take position 6, 7, and 8 in the onehot vector, and so on. This resulted in a high-dimensional sparse representation of the sequence. The collection date was converted to a numerical feature representing the number of days elapsed since the earliest collection date in the entire dataset (January 1, 1993). For the date prediction model, this numerical time feature was logtransformed (*y*^*′*^ = log(*y* + 1)) to stabilize variance and improve model performance, where *y* is days since the first sample. Geographic coordinates (latitude and longitude) were included as continuous numerical features.

### 3.2 Passage History Classification Model

#### 3.2.1 Model Architecture and Training

A LightGBM classifier [44, 45], a gradient boosting framework that uses tree-based learning algorithms, was trained to predict the passage history of influenza sequences. The input features for this model included the onehot encoded HA sequence, the numerical collection date (‘times’), latitude, and longitude. The single sequence belonging to the ‘VERO’ class was removed to ensure stable stratified splitting, resulting in a final dataset for passage classification comprising 39,121 sequences. This dataset was split into a training set (31,296 sequences, 80%) and a test set (7,824 sequences, 20%) using stratified sampling to maintain class proportions in both sets. The model was trained on the training set.

#### 3.2.2 Hyperparameter Optimization

Hyperparameters for the LightGBM classifier were optimized using a Bayesian optimization approach with the bayes opt Python library. The optimization aimed to maximize the 5-fold cross-validated balanced accuracy on the training set. The search space included parameters such as `num_leaves` (100–1000), `max_depth` (5–15), `n_estimators` (100–2000), `min_data_in_leaf` (2–30), and `reg_lambda` (100–1000). The optimization was run for 50 initialization points and 100 iterations. The best hyperparameters found were: `boosting_type=‘gbdt’`, `objective=‘multiclass’`, `metric=‘multi logloss’`, `learning rate=0.1`, `reg_alpha=0.0, reg lambda=522.0`, `min_data_in_leaf=5, n_estimators=1318`, `num_leaves=503, max_depth=8`, `class_weight=‘balanced’`, `colsample_bytree=1.0, n_jobs=-1`, and `random_state=42`. These optimized parameters were used for the final model training and evaluation.

#### 3.2.3 Model Evaluation

The performance of the passage classification model was evaluated on the held-out test set. Metrics included overall accuracy, balanced accuracy (to account for class imbalance), macro-averaged precision, recall, and F1-score. A normalized confusion matrix was generated to visualize the classification performance across different passage categories.

### 3.3 Sampling Date Prediction Model

#### 3.3.1 Dataset Preparation

For the task of predicting the sampling date, a separate LightGBM regressor was trained. This model used only sequences designated as ‘UNPASSAGED’ to avoid confounding signals from laboratory adaptation. This dataset comprised 20,109 unpassaged HA sequences. The data was split into a training set (16,087 sequences, 80%) and a test set (4,022 sequences, 20%) by random sampling.

#### 3.3.2 Model Architecture and Training

The LightGBM regressor was trained to predict the log-transformed collection date (days since the earliest sample). Input features for this model were restricted to the one-hot encoded HA sequence, latitude, and longitude. The collection date was not provided as an input feature to this model as it was the target.

#### 3.3.3 Hyperparameter Selection

Initially, I attempted Bayesian optimization as with the classifier. However, between runs there was substantial variability despite virtually all models being highly accurate and providing similar feature loadings. Thus, due to the regression task’s relative insensitivity to starting parameters observed in preliminary experiments, a fixed set of reasonable hyperparameters was used for the LightGBM regressor. These parameters were: boosting_type=‘gbdt’, objective=‘regression l2’, metric=‘rmse’, learning_rate=0.1, reg_alpha=0.0, reg_lambda=400.0, min_data_in_leaf=15, n_estimators=1200, num_leaves=215, max_depth=8, colsample_bytree=1.0, n_jobs=-1, and `random_state=42`.

#### 3.3.4 Model Evaluation

The performance of the date prediction model was evaluated on the test set using the coefficient of determination (R^2^), Mean Squared Error (MSE), and Mean Absolute Error (MAE). MAE was reported for both the log-transformed predictions and for predictions transformed back to the original scale of days. Predicted dates were plotted against actual collection dates to visually assess model fit. Residual plots (residuals vs. predicted values, and histogram of residuals) were also generated to check for systematic biases.

#### 3.3.5 Comparison with Simpler Models

To contextualize the performance of the full date prediction model, two simpler baseline models were trained and evaluated on the same unpassaged dataset splits: 1. A “sequence-only” LightGBM regressor using only the onehot encoded sequence features. 2. A “geo-only” LightGBM regressor using only latitude and longitude features. Both baseline models used the same hyperparameters as the full date prediction model. Their performance was assessed using R^2^ on the test set.

### 3.4 Model Interpretation using SHAP

To understand the contributions of individual features to the predictions of both the passage classifier and the date regressor, SHAP (SHapley Additive exPlanations) values were computed [42]. SHAP is a game theory-based approach that assigns an importance value to each feature for each prediction. I used the shap.TreeExplainer method, which is optimized for tree-based models like LightGBM.

For the passage classifier, SHAP values were calculated for each class. The mean absolute SHAP values across all test samples were used to rank features (amino acids per site) globally. These values were then aggregated by protein site (summing SHAP values for all amino acid possibilities at a site, and for non-sequence features like time, latitude, longitude) to determine overall site importance. For the stacked bar plot (Fig. 3), SHAP values were summed per protein site (or non-sequence feature). For visualization, the original shap value (whether positive or negative) was retained to demonstrate the feature or site’s impact on the model target.

For the date regressor, similar to above, mean absolute SHAP values were calculated for each feature across the test set. These were then aggregated by summing the importances for all one-hot encoded features corresponding to a single protein site to get an overall importance score for each of the 566 sites (in the final outputs the initial 16 sites were excluded as they are not in the mature protein). These site-level importances were then plotted. Again, the original SHAP value per amino acid per site (whether positive or negative) was retained to demonstrate the impact on the model target.

### 3.5 Sequence Variability and Evolutionary Rate Estimation

#### 3.5.1 Shannon Entropy

Shannon entropy was calculated for each of the 566 amino acid sites in the HA gene using the alignment of all 39,121 quality-filtered sequences. For each site, entropy *H*_*s*_ was computed as *H*_*s*_ = − Σ _*i*_ *p*_*i*_ log_2_ *p*_*i*_, where *p*_*i*_ is the frequency of amino acid *i* at that site.

#### 3.5.2 Relative Evolutionary Rate Estimation

Site-specific LEISR values for the HA protein were estimated using LEISR (Likelihood Estimation of Individual Site Rates) implemented in HyPhy [46]. This analysis was performed on the set of 20,109 unpassaged HA protein sequences. The input for LEISR was a codon alignment corresponding to these protein sequences and a phylogenetic tree. The LEISR method estimates the relative evolutionary rate for each protein site based on a given phylogenetic tree and substitution model. The output provided the Maximum Likelihood Estimate (MLE) of relative evolutionary rate for each gene site (1-566).

#### 3.5.3 Site-wise dN/dS Estimation

To estimate site-specific selection pressures, the ratio of nonsynonymous to synonymous substitution rates (dN/dS) was calculated using the FUBAR method (Fast, Unconstrained Bayesian AppRoximation) in HyPhy. As with the LEISR analysis, this was performed on the codon alignment of the 20,109 unpassaged sequences using the corresponding phylogenetic tree. FUBAR provides a robust and computationally efficient Bayesian approach to inferring site-wise selection. The posterior mean dN/dS values were used for downstream correlation analyses.

### 3.6 Phylogenetic Analysis and Root-to-Tip Regression

A phylogenetic tree was constructed for the 20,109 unpassaged HA sequences to perform root-to-tip distance calculations. The nucleotide alignment was inferred using IQ-TREE multicore version 2.3.1 [47] with the command `iqtree2 -s H3_nuc_unpassaged.fasta -st DNA -m GTR+G -fast -nt AUTO`. The general timereversible model with Gamma-distributed rate heterogeneity (GTR+G) was employed, as selected by ModelFinder [48]. The resulting tree was midpoint-rooted using the Bio.Phylo module from Biopython [49]. Root-to-tip distances (the sum of branch lengths from the root to each terminal node/tip) were then calculated for all sequences in the tree.

### 3.7 Biological and Structural Context

#### 3.7.1 Antigenic Site Annotation

To benchmark model-derived importance against known functional regions, I defined a set of seven experimentally validated antigenic sites (protein sites 145, 155, 156, 158, 159, 189, and 193) based on the work of Koel et al. [8]. These sites were mapped to my protein alignment, and this annotation was used to compare the performance of SHAP and classical metrics in prioritizing functionally critical residues.

#### 3.7.2 Structural Mapping and Visualization

To visualize the spatial location of important residues, site-level metrics were mapped onto the three-dimensional structure of the H3 HA protein. The crystal structure of A/Hong Kong/1/1968 HA in complex with a human receptor analog (PDB ID: 4O5N) was used as a reference. Structural visualization and figure generation were performed using The PyMOL Molecular Graphics System, Version 2.5 (Schrödinger, LLC). In Figure 10, the top ten SHAP-ranked sites from the passage model and the seven Koel et al. antigenic sites were highlighted on the protein backbone to illustrate their proximity to the sialic acid binding site.

### 3.8 Correlation Analyses

Spearman correlation coefficients (*ρ*) were calculated to assess the relationships between different metrics. For correlations involving SHAP values and entropy, relative evolutionary rates, or dN/dS, SHAP values (derived based on protein sites) were aggregated. Specifically, the mean absolute SHAP values for all features belonging to a given protein site were summed. These summed SHAP values per protein site were then mapped to the corresponding gene site (protein site + 16) to align with dN/dS, LEISR, and entropy data, which were indexed by gene site.

### 3.9 Temporal Amino Acid Frequency Analysis

To visualize evolutionary dynamics at key sites identified by the date prediction model, amino acid frequency trajectories were plotted over time. This analysis focused on the 20,109 unpassaged sequences. For selected protein sites with high SHAP importance in the date model, sequences were binned into 60-day intervals based on their collection dates. Within each time bin, the frequency of each amino acid at the selected site was calculated. These frequencies were then plotted as a time series to illustrate changes in amino acid prevalence.

### 3.10 Software and Data Availability

Analyses were performed using Python (version 3.x). Key Python libraries included LightGBM for gradient boosting models, SHAP for model interpretation, scikit-learn for machine learning utilities (e.g., train-test splitting, metrics, OneHotEncoder), pandas for data manipulation, NumPy for numerical operations, Biopython for sequence and phylogenetic tree handling, flupan for passage history parsing, Matplotlib and Seaborn for plotting, geopy for geocoding, and bayes opt for Bayesian hyperparameter optimization. Phylogenetic tree inference was performed with IQ-TREE (version 2.3.1). Relative evolutionary rate estimation was performed using LEISR and FUBAR from the HyPhy package. All HA sequences and associated metadata used in this study were sourced from GISAID (www.gisaid.org), and I grateful for the contributions of the originating laboratories and submitting laboratories for making these data available. Specific GISAID accession numbers and the code used for analysis in this study is available at https://github.com/ausmeyer/flulightgbm shap submission.

